# Task-Switch Related Reductions in Neural Distinctiveness in Children and Adults: Commonalities and Differences

**DOI:** 10.1101/2023.12.22.572358

**Authors:** Sina A. Schwarze, Sara Bonati, Radoslaw M. Cichy, Ulman Lindenberger, Silvia A. Bunge, Yana Fandakova

## Abstract

Goal-directed behavior requires the ability to flexibly switch between task sets with changing environmental demands. Switching between tasks generally comes at the cost of slower and less accurate responses. Compared to adults, children show greater switch costs, presumably reflecting the protracted development of the ability to flexibly update task-set representations. To examine whether the distinctiveness of neural task-set representations is more strongly affected by a task switch in children compared to adults, we examined multi-voxel patterns of fMRI activation in 88 children (8–11 years, 49 girls) and 53 adults (20–30 years, 28 women) during a task-switching paradigm. Using multivariate pattern analysis (MVPA), we investigated whether task-set representations were less distinct on switch than on repeat trials across frontoparietal, cingulo-opercular, and temporo-occipital regions. Children and adults showed lower accuracy and longer response times on switch than on repeat trials, with higher accuracy costs in children. Decoding accuracy across regions was lower on switch than repeat trials, consistent with the notion that switching reduces the distinctiveness of task-set representations. Reliable age differences in switch-related representational distinctiveness reductions were absent, pointing to a remarkable degree of maturity of neural representations of task-relevant information in late childhood. However, we also observed that switch-related reductions in distinctiveness were more highly correlated across frontoparietal and cingulo-opercular regions in children than in adults, potentially reflecting the ongoing specialization of different control networks with respect to the representation of task features.

**Significance statement:** The ability to flexibly switch between tasks enables goal-directed behavior, but is particularly challenging for children, potentially due to protracted development in the ability to represent multiple and overlapping task rules that link stimuli to appropriate responses. We tested this hypothesis by using neuroimaging to measure brain activity during task switching in 8–11-year-olds and adults. Activation patterns in frontal, parietal, and temporal regions could tell us with above-chance accuracy which task a person was performing when the task remained the same, but not when it had switched. Adults showed greater differentiation across regions in terms of switch-related reductions in distinctiveness than children, suggesting that the relevant functional circuity is present but has not yet fully matured by late childhood.

## 1. Introduction

The ability to flexibly switch between tasks, thoughts, or actions when circumstances change is critical for goal-directed behavior (Miyake and Friedman, 2012; Diamond, 2013). However, switching between tasks entails a cost compared to repeating previously executed tasks, such that responses are slower, less accurate, or both. A task switch requires updating the task set, which amounts to retrieving the task set for the newly relevant task and inhibiting the no-longer relevant task set (Rogers and Monsell, 1995; Meiran, 1996; Mayr and Kliegl, 2000; Wylie and Allport, 2000; for a review, see Vandierendonck et al., 2010). Both of these processes are thought to contribute to the switch costs observed in performance.

Behaviorally, switch costs are more pronounced in children compared to adults (Crone et al., 2006a; Huizinga et al., 2006; Huizinga and van der Molen, 2007; Gupta et al., 2009; Cragg and Chevalier, 2012; Church et al., 2017; but see Luca et al., 2003; Reimers and Maylor, 2005). Age-related differences in switch costs have been attributed to children’s difficulties to inhibit the no-longer relevant task set and to update the relevant task set when rules switch (Crone et al., 2004, 2006a; Gupta et al., 2009; Wendelken et al., 2012). Moreover, children’s representations of goal-relevant task sets have been suggested to be less distinct from one another (Zelazo, 2004; Crone et al., 2006b; Lorsbach and Reimer, 2008; Jung et al., 2023), especially when task sets are partially overlapping (e.g., due to same responses; cf. Crone et al., 2006a).

Children’s task-switching difficulties have been associated with smaller increases in activation for switch compared to repeat trials in frontoparietal (FP) brain regions, including the inferior frontal junction (IFJ), the superior parietal lobe (SPL), and the dorsolateral prefrontal cortex (dlPFC), compared to adults (Crone et al., 2006b; Bunge and Wright, 2007; Velanova et al., 2008; Wendelken et al., 2012; Engelhardt et al., 2019; Zhang et al., 2021; Schwarze et al., 2023; but see Morton et al., 2009). The previous studies did not examine differences in multivariate patterns of neural activations between repeat and switch trials, and therefore do not address whether age differences therein are a possible source of age-related differences in switch costs.

Research in adults (Loose et al., 2017; Qiao et al., 2017) has started to feature multivariate pattern analysis (MVPA; Haynes and Rees, 2006) to examine the distinctiveness of neural representations of task sets in task-switching paradigms. Because task-set representations on switch trials have just been updated, they are hypothesized to be less distinct on switch compared to repeat trials (Meiran, 1996; Mayr and Kliegl, 2000), resulting in lower decoding accuracy. Decoding accuracy describes how well the currently relevant task can be predicted from the pattern of neural activation. Beyond this updating process, the lingering representation of the previously relevant task set (i.e., task-set inertia) might dilute the current task-set representation, further contributing to less distinct representations on switch compared to repeat trials (Rogers and Monsell, 1995; Wylie and Allport, 2000; Rangel et al., 2023). Studies investigating this hypothesis in adults have reported contradictory results. While one study showed greater decoding accuracy on repeat than on switch trials (Qiao et al., 2017), other studies showed the opposite pattern (Tsumura et al., 2021) or no differences between conditions (Loose et al., 2017). To date, neural task-set representations have not been examined in children to provide a direct test of representational accounts of children’s difficulties in task switching.

We used MVPA to assess the distinctiveness of task-set representations in children (N = 88, 8–11 years) and adults (N = 53, 20–30 years) who performed a task-switching paradigm during neuroimaging. We expected that, children would show overall lower decoding accuracy and would be disproportionally affected by the demand to switch, resulting in lower decoding accuracy on switch trials compared to adults. To explore whether age differences in task switching were related to stronger task-set inertia in children than in adults (Gupta et al., 2009; Hommel et al., 2011; Wendelken et al., 2012; Witt and Stevens, 2012; but see Crone et al., 2006a), we used three different task sets. As opposed to switches among two task sets, this design allowed us to test whether decoding accuracy for the previously relevant task on switch trials was higher compared to the third task (that was neither relevant on the current nor on the previous trial), which would indicate task-set inertia.

While adult studies of neural representations during task switching have focused on FP regions, the distinctiveness of task-set representations in temporo-occipital (TO) regions (i.e., fusiform gyrus, parahippocampal gyrus, and lateral occipital cortex) may be particularly important for task switching in children. These regions mature earlier than FP regions do (Sydnor et al., 2021) and may contribute to the development of more distinct FP representations during rule-based tasks (Amso and Scerif, 2015; Rosen et al., 2019). Furthermore, regions of the cingulo-opercular (CO) network, including the dorsal anterior cingulate cortex (dACC) and the anterior insula (aI), have been associated with task-set maintenance during switching (Braver et al., 2003; Dosenbach et al., 2008; Gratton et al., 2018). Due to the relatively more sustained nature of maintenance processes (Braver et al., 2003), representations in CO regions may not be updated in a trial-specific fashion, resulting in smaller differences in decoding accuracy between switch and repeat trials. Thus, we explored how the distinctiveness of task-set representations in brain regions associated with different roles during task switching (i.e., FP vs. CO) and with different developmental trajectories (i.e., FP/CO vs. TO regions) differed with respect to age differences.

## 2. Materials and Methods

The hypotheses and the analysis plan were preregistered before the start of analysis (https://osf.io/8mfqx/). We explicitly note deviations from the preregistered analysis plan below. The behavioral performance and univariate analysis of functional neuroimaging data of this sample are described in detail in Schwarze et al. (2023), and are only briefly summarized below.

### 2.1 Participants

Children (N = 117) and adults (N = 53) completed the task-switching paradigm in the MR scanner, as reported previously (Schwarze et al., 2023). All participants were right-handed. Prior to running the analyses of interest, participants were excluded if they showed low accuracy (i.e., below 50% in the fMRI run of single tasking or below 35% in either of the two runs that presented tasks intermixed, see details below; N = 8 children excluded), excessive in-scanner motion (more than 50% of fMRI volumes with framewise displacement (Power et al., 2012) above 0.4 mm; N = 24 children, 4 of whom also showed poor performance), or fewer than 5 trials for each class, i.e., each combination of condition (switch vs. repeat) and task (face vs. scene vs. object, see below), in the MVPA analyses (N = 1 child; included in the previous study [Schwarze et al., 2023]). The final sample included 88 children (8–11 years; mean age = 10.07 years, SD = 0.69; 49 girls) and 53 adults (20–30 years; mean age = 24.69 years, SD = 2.6; 28 women). Adult participants, parents, and children provided informed written consent and the study was approved by the ethics committee of the Freie Universität Berlin and conducted in line with the Declaration of Helsinki.

### 2.2 Experimental design

In the task-switching paradigm performed in the MR scanner, participants had to respond to one of three simultaneously presented stimuli (a face, a scene, and an object) each relevant to a different task (i.e., the face task, the scene task, and the object task). The relevant task was indicated by a simultaneously presented shape in the background, such that, for instance, a diamond-shaped background indicated the face task, which required the classification of the face according to its age (see Figure 1A). On each trial, the spatial arrangement of stimuli varied pseudo-randomly independent of the currently relevant stimulus. Participants responded via button press.

**Figure 1:**
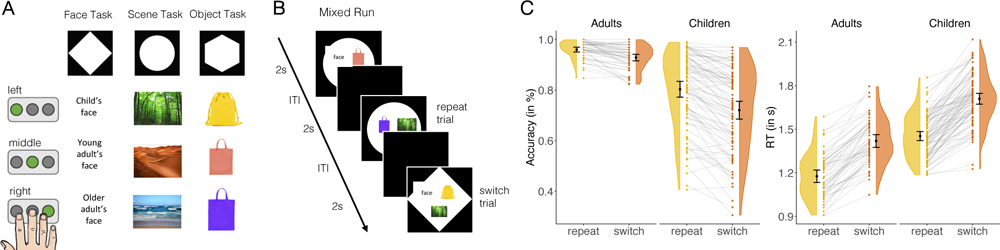
Task-switching paradigm and age differences in performance. (A) The task-switching paradigm consisted of three tasks: the face task, the scene task, and the object task. Participants had to perform the task indicated by the simultaneously presented shape in the background. Depending on the stimulus presented, one of three buttons had to be pressed in response (indicated here by the green button). Faces had to be categorized according to the age of the person shown, scenes according to the location (i.e., forest, desert, or sea), and objects according to the color. (B) The mixed runs included trials on which the task of the previous trial was repeated (50%) or switched to a different task (50%) in an unpredictable manner. (C) Performance accuracy (in %) and response times (in seconds) for repeat (yellow) and switch (orange) trials split by age group. Gray lines connect performance measures for each individual. Image credits: Young and old adult faces were taken from the FACES collection (Ebner et al., 2010).

The fMRI session consisted of three runs: one single run in which the tasks were presented in separate blocks, followed by two mixed runs in which the three tasks were pseudo-randomly intermixed. Each run had 99 trials each lasting 2s, followed by a fixation cross for a jittered time period (1–6 s). In the single run, tasks were presented in three separate blocks of 33 trials, interspersed with blocked fixation cross periods (20 s). In the mixed runs, the three tasks were pseudo-randomly intermixed, with 50% repeat and 50% switch trials, again across blocks of 33 trials with blocked fixation cross periods (20 s) to match the single run. Switches were unpredictable, such that participants did not know which task had to be performed on the upcoming trial. The first trial of each run was excluded from all analyses as it could not be classified as a switch or repeat trial in the mixed runs. The main MVPA determining representational changes during task switching focused on the two mixed runs. Data from all three runs were used to define regions of interest (ROIs) that were representative of the task-related univariate activation.

### 2.3 Behavioral measures and analysis

Behavioral results were previously reported for essentially the same sample across all three task runs (89 as opposed to 88 children, cf. Schwarze et al., 2023), whereas the present analysis focused on the two mixed runs and thus only on repeat and on switch trials. Individual trials with response times (RT) below 0.2 s or above 3 s were excluded from all analyses, so that responses during stimulus presentation and within the first second of the inter-trial interval were accepted. Only correct trials were considered for the calculation of median RTs per condition. Accuracy was calculated as the percentage of correct responses across all given responses for each condition. No datapoints of individual participants had to be removed based on the predefined outlier criterion of mean accuracy or median RTs that deviated 3.5 standard deviations (p < .001) or more from the age group-specific mean (Tabachnick and Fidell, 2013). We used Bayesian linear mixed models, using brms (version 2.19.0; Bürkner, 2017) in R (version 4.3.1; R Core Team, 2018), with flat priors to predict proportions of correct responses or median correct RT from condition, age group, and their interaction, including random intercepts for subject. For all models, reported effects are based on 95% credible intervals (CI) such that the described effects have a 95% probability in the present data (Bürkner, 2017; see also Morey et al., 2016).

### 2.4 fMRI data acquisition and preprocessing

Functional MR images were collected on a 3-Tesla Siemens Tim Trio MRI scanner using whole-brain echo-planar images (TR = 2000ms; TE = 30ms; 3 mm isotropic voxels). The first five acquired volumes of each run were discarded before analysis to allow for scanner stabilization.

Preprocessing was performed using fMRIprep (Version 20.2.0; Esteban et al., 2019; for a detailed description of procedures, see https://fmriprep.org/en/stable/). BOLD images were co-registered to individual anatomical templates using FreeSurfer, which implements boundary-based registration (Greve and Fischl, 2009). Additionally, they were slice-time corrected (using AFNI; Cox and Hyde, 1997), and realigned (using FSL 5.0.9; Jenkinson et al., 2002). For the definition of ROIs based on group-level univariate activation, BOLD images were normalized into MNI152NLin6Asym standard space. All multivariate analyses were conducted in individual-specific anatomical space. ROIs defined in MNI space were transformed into individual-specific anatomical space using Advanced Normalization Tools (ANTs; Avants et al., 2009) and FSL (Version 5.0.9; Jenkinson et al., 2002).

### 2.5 ROI definition

*Frontoparietal (FP) and cingulo-opercular (CO) regions.* Previous research has demonstrated that lateral FP regions, including the IFJ, the SPL, and dlPFC, show enhanced univariate activation during task switching (Kim et al., 2012; Niendam et al., 2012; Richter and Yeung, 2014; Worringer et al., 2019) and represent the currently relevant task in adult studies (Loose et al., 2017; Qiao et al., 2017). Thus, we focused on these areas as our main ROIs (see preregistration: https://osf.io/8mfqx/). Along with the expected FP regions, the univariate activation analyses (see Figure 2) also revealed enhanced activation in the dACC and the aI during task switching. Thus, for exploratory analyses, we defined ROIs of the CO network, including the aI and dACC (Dosenbach et al., 2007, 2008).

**Figure 2:**
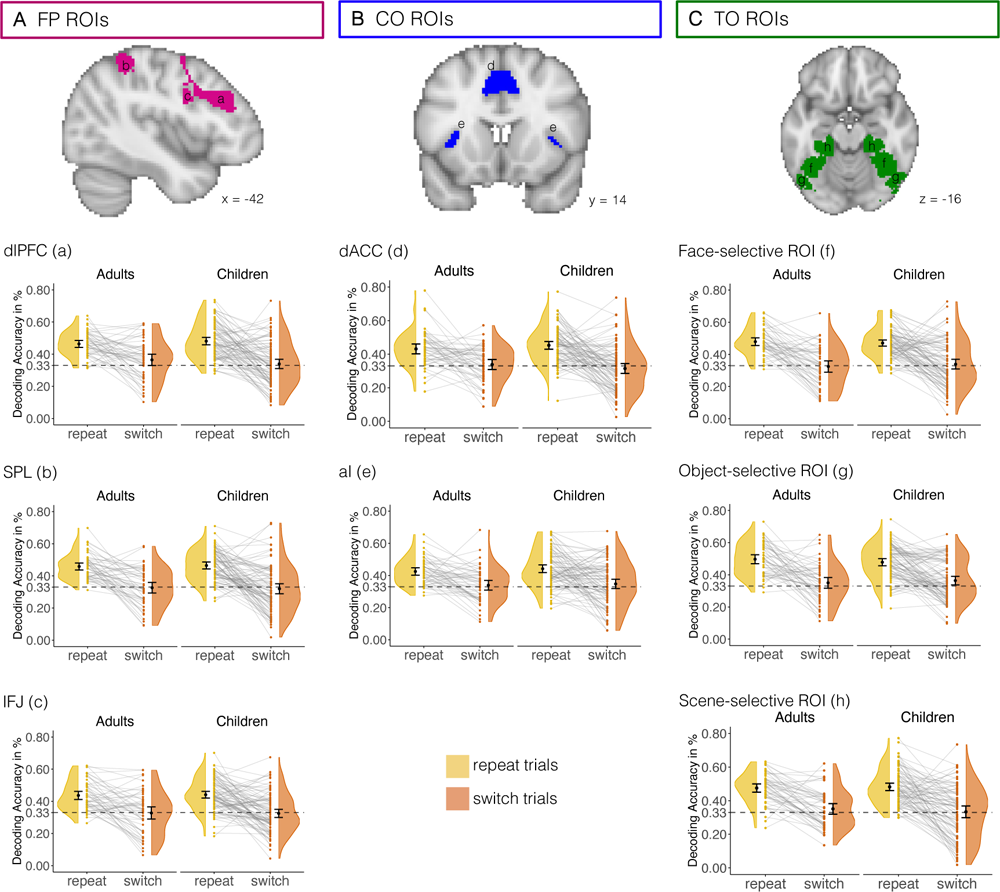
Regions of interest (ROIs) and decoding accuracy results. The dashed line in each plot indicates chance (0.33). (A) Decoding accuracy for repeat trials (yellow) and switch trials (orange) in adults and children of frontoparietal (FP) regions: (a) dorsolateral prefrontal cortex (dlPFC), (b) superior parietal lobe (SPL), and (c) inferior frontal junction (IFJ). (B) Decoding accuracy of cingulo-opercular (CO) regions: (d) dorsal anterior cingulate cortex (dACC) and (e) anterior insula (aI). (C) Decoding accuracy of temporo-occipital (TO) regions: (f) face-selective ROI in the fusiform gyrus, (g) object-selective ROI in the lateral occipital cortex, (h) scene-selective ROI in the parahippocampal gyrus.

The ROI definition procedure was as follows. ROIs were initially defined based on task activation across all three runs relative to baseline and subsequently restricted by anatomical location. To this end, we constructed a general linear model (GLM) of correct single, correct repeat, and correct switch trials as separate regressors. Missed trials, error trials, and the first trial of each run were included in a separate regressor of no interest. Framewise displacement per volume (in mm; Power et al., 2012), realignment parameters (three translation and three rotation parameters), and the first six anatomical CompCor components as provided by fMRIprep were added as regressors of no interest. CompCor identifies patterns of noise using a principle-component analysis approach and the inclusion of the components aids in the removal of noise from fMRI data (Behzadi et al., 2007). We derived a contrast, comparing task (correct single, repeat, and switch trials) with baseline collapsed across age groups. The resulting whole-brain contrast map was thresholded at family-wise error (FWE) corrected p < .05, cluster size > 50 voxels.

Multiple brain regions in the frontal and parietal cortices showed greater activation for tasks compared to baseline, including bilateral IFJ, dlPFC, SPL, dACC, and aI. Functional activations as determined above were anatomically restricted using the Harvard-Oxford atlas (Makris et al., 2006), thresholded at 30%. The inclusive anatomical masks we used to restrict univariate activation to pre-defined ROIs were the middle frontal gyrus for the dlPFC, the superior parietal lobe for the SPL, insular cortex for the aI, and the paracingulate gyrus for the dACC. Because no anatomical mask for the restriction of the IFJ is available, we defined it based on coordinates from a meta-analysis of task-switching studies focusing on the IFJ (Derrfuss et al., 2005). See Figure 2 A–B for task-based FP and CO ROIs, respectively.

*Temporo-occipital (TO) ROIs*. ROIs in TO were defined on activation maps provided by the Neuroquery (Dockès et al., 2020) and Neurosynth (Yarkoni et al., 2011) platforms, using the search terms “face” “object”, and “place”. A probability map was downloaded for each of the search terms from each platform on November 22nd, 2021. Neuroquery maps were thresholded with a z-score of 3 as recommended by the developers (Dockès et al., 2020); Neurosynth maps were thresholded at p < .01 (FDR-corrected; Yarkoni et al., 2011). All negative value voxels were set to zero to keep only positive activation associated with each search term. Next, the Neurosynth and Neuroquery masks for each search term were multiplied with each other to only include voxels identified across both platforms. Note that for the “place” search term only the Neuroquery mask was used in ROI definition, as the map provided by Neurosynth did not include the parahippocampal gyrus, which has consistently been associated with place/scene perception across age groups (Golarai et al., 2007; Scherf et al., 2007).

Finally, following the same approach as for the FP and CO ROIs, the resulting maps were anatomically masked using the Harvard-Oxford atlas (Makris et al., 2006), thresholded at 30%. The temporal occipital fusiform gyrus was used as an anatomical mask for the face-selective ROI, the inferior lateral occipital cortex for the object-selective ROI, and the anterior and posterior parahippocampal gyrus for the scene/place-selective ROI. The resulting TO ROIs overlapped with activation in temporo-occipital regions of the task > baseline contrast described above. Figure 2C shows the resulting TO ROIs.

Note that we had originally preregistered to define ROIs based on a searchlight MVPA decoding the three tasks (face vs. scene vs. object) across all runs, but we changed our approach due to updated methods for applying corrections of unequal class counts (see below).

### 2.6 Multivariate pattern analysis

We constructed subject-specific GLMs of the task-switching paradigm using the Nipype (version 1.6.0; Gorgolewski et al., 2011) interface to FSL FEAT (using FSL 5.0.9; Jenkinson et al., 2002), focusing on the mixed runs and more specifically, on the differences between switch and repeat trials therein. GLMs included the combination of task and condition as separate regressors (i.e., face switch, face repeat, scene switch, scene repeat, object switch, object repeat) and the same nuisance regressors of no interest as in the GLM for ROI definition. Activation patterns for individual trials in the two mixed runs were extracted using a least squares separate approach, in which a trial-specific design matrix is used to obtain the activation estimate for that trial (LSS; Mumford et al., 2012).

Next, we conducted MVPA for each participant in each condition separately, using Nilearn (version 0.8.0; see Abraham et al., 2014) and scikit-learn (version 0.24.2; Pedregosa et al., 2011). We used a support vector classifier (LinearSVC, initialized with regularization parameter C = 1 and one-vs-rest multiclass strategy) trained to predict the currently relevant task (scene, object, face), given trial activation patterns in each ROI. We applied leave-one-run-out cross-validation during the analysis, so that at each validation fold, a classifier was fitted on data from one run and tested on data of the other run. Participants with fewer than five trials of each task in a condition were excluded from analysis. To ensure balanced numbers of trials across classes, we undersampled the majority class(es) without replacement. We adopted this approach as opposed to the preregistered method of the scikit-learn “balanced” option (Pedregosa et al., 2011) as it eliminates possible bias rather than correcting for it post-hoc. We repeated model training and testing with leave-one-group-out cross-validation 100 times for each participant in each condition and averaged across iterations. We did not have to remove any participant’s decoding accuracy data based on the predefined outlier criterion of 3.5 standard deviations (p < .001) above or below the group-specific mean.

### 2.7 Analysis of age differences in decoding accuracy

#### 2.7.1 Age and condition differences within ROI sets

To test whether decoding accuracy differed between switch and repeat trials and between the two age groups, analyses in the three sets of ROIs (FP regions: dlPFC, IFJ, SPL; CO regions: dACC, aI; TO regions: fusiform gyrus [face-selective], parahippocampal gyrus [scene-selective], and lateral occipital cortex [object-selective]) proceeded in three steps: (1) For each ROI, we tested whether it would be appropriate to combine data across hemispheres: that is, we tested whether decoding accuracy differed between hemispheres, and whether hemisphere interacted with age group or condition. We did not find main effects or interactions involving hemisphere in any of the ROIs and thus averaged across the two hemispheres for all subsequent analyses. (2) Within each ROI set (FP, CO, TO), we tested a Bayesian linear mixed model across all regions in the corresponding set. Decoding accuracy was predicted by condition (repeat vs. switch), ROI, age group (adults vs. children), and their interactions. All linear mixed models included a random intercept for participant. If an effect of region or any interaction of region with another effect of interest (condition or age group) became evident with 95% probability (i.e., the 95% CI did not include zero), we proceeded to test for the main effects and interactions of condition and age group in each ROI separately. (3) Using t-tests, we tested whether decoding accuracy for each condition in each age group and ROI differed from chance (i.e., 0.33). In addition to comparing decoding accuracy between switch and repeat trials across all three tasks, we tested whether each of the TO ROIs showed selectivity for the theoretically preferred task on repeat trials. To this end, we compared repeat decoding accuracy of the preferred task (i.e., the face task for the fusiform gyrus) to the two non-preferred tasks (i.e., the scene and object task for the fusiform gyrus), and whether this effect differed between age groups.

#### 2.7.2 Age and condition differences between ROI sets

To examine whether age group and condition effects differed between the three sets of ROIs, we first modeled decoding accuracy including all ROIs. Specifically, models included fixed effects of condition, age group, and set of ROIs (FP vs. CON vs. TO), and their interactions, along with random slopes for set of ROI and random intercepts for participant. The model including all interactions fit slightly better than a model without any interactions of condition.

#### 2.7.3 Task-set inertia

In a set of exploratory analyses, we explored if the lingering representation of the previously relevant task set contributed to lower decoding accuracy on switch trials. Specifically, we tested whether incorrect predictions of the classifier on switch trials were more likely to predict the previously relevant task. As described above, the classifier predicted one of the three tasks for each trial. This prediction could either be correct and thus count towards the decoding accuracy measure, or incorrect if the prediction indicated one of the other two tasks not relevant on that specific trial. To test the task-set inertia hypothesis within each set of ROIs, we tested whether the classifier was more likely to predict the previously relevant task over the task that was neither relevant on the previous nor current trial; further, we tested whether this differed between age groups. To this end, we modelled percentage of false predictions as the dependent variable and the type of false prediction (previous vs. third task) and age group as the fixed effects, additionally including fixed effects of region and random intercepts of participant.

#### 2.7.4 Individual differences in the impact of switch demand on representations across ROI sets

While analyses up to this point tested whether the effect of switch demand on neural task-set representations differed between children and adults, they did not shed light on the question whether switch demand affected different brain regions similarly within an individual. Thus, to further understand these individual differences across the sets of ROIs, we explored whether individuals who showed a greater difference in decoding accuracy between conditions in one set of ROIs showed a similar pattern in the other sets of ROIs. To this end, we averaged the differences between switch and repeat decoding accuracy across all ROIs in each set. Next, we tested for age group differences in the correlations among the three sets of ROIs by comparing the correlations of each pair of ROI sets between the two age groups using cocor (version 1.1-4; Diedenhofen and Musch, 2015) in R.

### 2.8 Associations between decoding accuracy and performance

To anticipate the outcome of our analyses, we did not observe any differences in decoding accuracy between individual ROIs within each set. As a result, we deviated from the preregistration and tested whether decoding accuracy across ROIs in a set predicted task performance. We used a linear mixed model with performance accuracy as the dependent variable, average decoding accuracy across the ROIs in one set, condition (repeat vs. switch), and age group (adults vs. children) as fixed effects, and a random intercept modeling the individual participants. We used leave-one-out cross-validation (loo package; Vehtari et al., 2022) to compare the model including all interactions between the fixed effects to models including fewer interaction terms. We only tested models that included an interaction of age group and condition to account for differences in behavioral performance between children and adults. For both behavioral accuracy and RT, and the models in all ROI sets, model comparisons indicated better fit for the model including only the main effect of decoding accuracy and the interaction between age group and condition, but no interaction of decoding accuracy with either age group or condition. Thus, the effect of interest was the main effect of decoding accuracy. The same model setup and comparison approach was used for linear mixed models of RT.

## 3. Results

### 3.1 Greater switch costs in children than adults

Adults exhibited higher overall performance accuracy than children (estimate (est.) = –0.16; 95%-CI: –0.20, –0.11) as well as shorter RTs on correct trials (est. = 0.28; 95%-CI: 0.22, 0.33). Both groups exhibited switch costs, with higher accuracy on repeat than on switch trials (est. = –0.03; 95%-CI: –0.04, –0.02); as well as shorter correct RTs on repeat than on switch trials (est. = 0.24; 95%-CI: 0.21, 0.27). Critically, children exhibited greater switch costs than adults in terms of accuracy (condition x group interaction: est. = –0.05; 95%-CI: –0.07, –0.03; Figure 1C), albeit not in terms of RTs (condition x group interaction: est. = 0.01; 95%-CI: –0.03, 0.05; Figure 1C). In sum, both children and adults showed switch costs in accuracy and RT, with greater accuracy switch costs in children than in adults.

### 3.2. Higher decoding accuracy for repeat than for switch trials across age groups

Decoding accuracy for each ROI is shown in Figure 2. We predicted lower decoding accuracy on switch than on repeat trials across groups in all sets of ROIs, with greater reductions in decoding accuracy for children. To test these hypotheses, we used Bayesian linear mixed models for each set of ROIs to predict decoding accuracy by age group, condition, and ROI.

In each set of ROIs, we found evidence of a main effect of condition (switch vs. repeat; FP ROIs: est. = –0.10; 95%-CI: –0.14, –0.06; CO ROIs: est.= –0.09; 95%-CI: –0.13, –0.05; TO ROIs: est. = –0.15; 95%-CI: –0.19, –0.11), with higher decoding accuracy on repeat than on switch trials. There was no evidence for effects of age group in any of the ROI sets, showing that, contrary to our hypothesis, decoding accuracy was comparable between children and adults. Finally, there were no effects of specific ROI within a set, suggesting that all tested ROIs showed higher decoding accuracy on repeat trials relative to switch trials in both age groups.

ROI-specific analyses indicated that all ROIs showed above-chance (> 0.33) decoding accuracy on repeat trials (all ts > 6.85; all lower bounds of 95%-CI > 0; see Figure 2). By contrast, decoding accuracy on switch trials did not differ from chance in the majority of ROIs (i.e., the bilateral SPL and IFJ for the FP ROIs, the bilateral aI and dACC of the CO ROIs, and the face-and scene-selective ROI; –1.04 < t < 1.36; all lower bounds of 95%-CI < 0), with only two exceptions. The dlPFC showed above-chance decoding accuracy for switch trials in adults (t = 1.96; lower bound of 95%-CI = 0.005) but not in children (t = 0.64; lower bound of 95%-CI = –0.015), and the object-selective ROI showed above-chance decoding accuracy for switch trials in children (t = 2.55; lower bound of 95%-CI = 0.012), but not in adults (t = 1.2; lower bound of 95%-CI = –0.0077).

Taken together, in line with our hypothesis and in accordance to the observed behavioral switch costs, decoding accuracy was greater for repeat than for switch trials across all sets of ROIs. Above-chance decoding of the currently relevant task was only evident for repeat trials, while the newly-updated relevant task on a switch trial could not be distinguished from the two irrelevant tasks based on the neural activation pattern. Of note, contrary to our hypothesis, children showed comparably distinct task-set representations as adults.

To explore whether the TO ROIs showed preference for stimuli in the currently relevant task, given their putative functional specialization for processing different kinds of stimuli (Cantlon et al., 2011; Natu et al., 2016; Golarai et al., 2017; Tian et al., 2021), we tested decoding accuracy for the preferred compared to the non-preferred tasks on repeat trials in each ROI. The face-selective ROI showed greater decoding accuracy on repeat trials for the face task compared to the object and scene tasks (est. = –0.06; 95%-CI: –0.8, –0.03) even though each trial presented a face, object, and scene stimulus simultaneously on the screen and at unpredictable locations. By contrast, neither the object-nor scene-selective ROI showed a preference for object or scene tasks, respectively. Thus, only the face-selective ROI showed a preference for the stimuli it was expected to prefer.

### 3.3 Higher decoding accuracy for temporo-occipital ROIs

As the effect of condition was present in all three sets of ROIs, we next sought to directly compare whether it differed between the sets of ROIs. A model comparing the effects of age group, condition, and ROI set on decoding accuracy revealed that relative to the TO ROIs, decoding accuracy was lower in the FP ROIs (est. = –0.03; 95%-CI: –0.06, –0.01) and CO ROIs (est. = –0.06; 95%-CI: –0.06, –0.04). The difference between the CO and the TO ROIs was further qualified by an interaction with condition (est. = 0.06; 95%-CI: 0.02, 0.1), indicating a greater difference in decoding accuracy between switch and repeat trials in the TO compared to the CO ROIs. There was neither evidence for differences between the FP and the CO ROIs nor for any differences between the age groups. Thus, while all investigated regions showed greater decoding accuracy on task repetitions compared to task switches, this condition difference, as well as overall decoding accuracy, was greater in the TO ROIs than in regions classically associated with cognitive control processes in children and adults.

### 3.4 No evidence for task-set inertia effects on representations

Next, we investigated whether task-set inertia contributed to less distinct task-set representations on switch trials. To this end, we compared the percentage of trials on which the classifier falsely predicted the previously relevant task to the percentage of trials on which it predicted the third task that was neither relevant on the current nor on the previous trial. Separate models for each set of ROIs included the fixed effects of incorrectly predicted task (previous vs. third task), age group, and region. None of the investigated sets of ROIs showed a higher probability of predicting the previous task over the third task in both children and adults (all 95%-CI included zero; Figure 3). Thus, we did not find any evidence that lower decoding accuracy on switch trials was related to task-set inertia, whereby the representation of the task relevant on the immediately preceding trial would linger after ceasing to be relevant.

**Figure 3:**
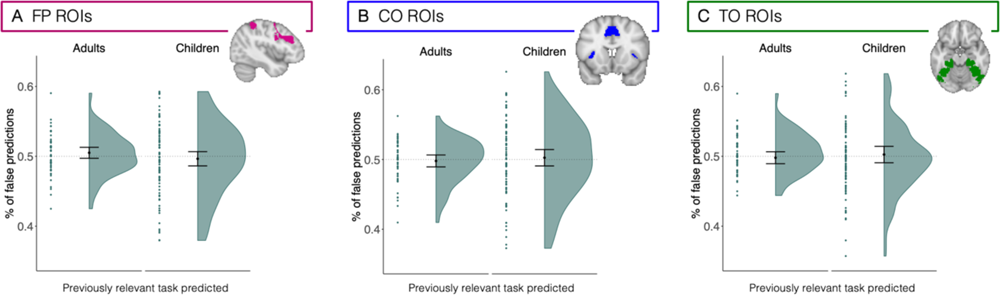
Percentage of predictions of the previously relevant task of all false predictions of the currently relevant task (i.e., task-set inertia) for (A) the frontoparietal (FP) ROIs, (B) the cingulo-opercular (CO) ROIs, and (C) the temporo-occipital (TO) ROIs. As none of the analyses within each set of ROIs indicated differences between regions, data were averaged across ROIs in each set for visualization.

### 3.5 Children showed similar condition differences in decoding accuracy in CO and FP ROIs

Next, to assess whether switch demand affected different brain regions similarly within an individual, we conducted a set of exploratory analyses testing whether children and adults showed similar patterns of switch-related reductions in decoding accuracy across ROI sets. To this end, we calculated the average difference in decoding accuracy between switch and repeat trials (i.e., difference scores) across all ROIs of the same set. The association between difference scores in the FP and in the CO ROIs differed between the age groups (p = .018; Figure 4): children showed a moderate to strong correlation between decoding accuracy difference scores in FP and CO (r = .64; p_FDR_ < .001, FDR-corrected for multiple comparisons), whereas adults showed a weak to moderate correlation (r = .33; p_FDR_ = .027). Difference scores showed a moderate correlation between FP and TO ROIs in children (r = .54; p_FDR_ < .001) and in adults (r = .38; p_FDR_ = .015), and did not differ between age groups (p = .27). Difference scores between CO and TO ROIs showed weak to moderate correlations in both children (r = .31; p_FDR_ = .004) and adults (r = .30; p_FDR_ = .03), and did not differ between age groups (p = .98). In sum, children showed greater similarity than adults in the impact of switching demands on FP and CO ROIs, while group differences were neither evident for the correlations between FP and TO nor for those between CO and TO.

**Figure 4:**
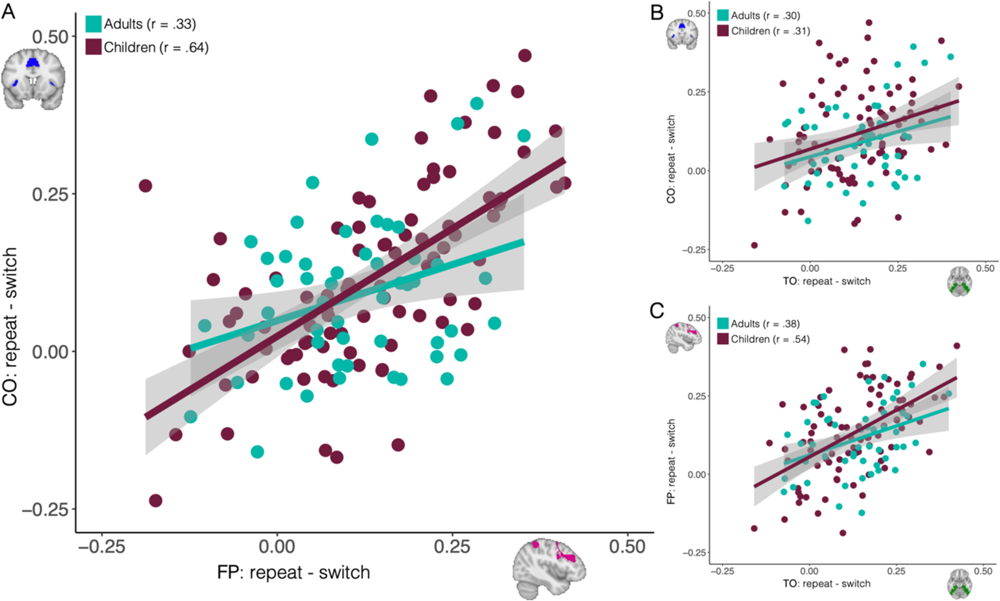
Correlations of condition differences in decoding accuracy between sets of ROIs split by age group. (A) In children (shown in magenta), greater differences in decoding accuracy between repeat and switch trials in frontoparietal (FP) ROIs were associated with greater differences on the same measure in the cingulo-opercular (CO) ROIs. This was not the case in adults (shown in turquoise). (B) Correlations in CO and temporo-occipital (TO) ROIs did not differ between children and adults. (C) Correlations in FP and TO ROIs did not differ between children and adults.

### 3.5 Decoding accuracy was not related to task-switching performance

Finally, to investigate whether having more distinct neural representations of the currently relevant task set was related to better task-switching performance, we tested whether decoding accuracy averaged across all ROIs within each set was associated with higher performance accuracy and/or lower RTs. Model comparisons indicated that the best fitting model included the main effect of decoding accuracy and the interaction between age group and condition, but no interaction of decoding accuracy with either age group or condition. This suggests that there were no differences between children and adults in the potential association between decoding accuracy and performance. In these models across all participants, none of the ROI sets showed an effect of decoding accuracy on either performance accuracy or correct RTs (all 95%-CI included zero). Thus, we found no evidence that higher decoding accuracy was associated with better performance during task switching in the present paradigm.

## 4. Discussion

Using MVPA, we examined the extent to which the distinctiveness of neural task-set representations contributed to age differences in task switching. Both children and adults showed lower decoding accuracy on switch compared to repeat trials across FP, CO, and TO regions, suggesting less distinct task-set representations on switch trials. We observed above-chance classification of the currently relevant task for repeat trials in all ROIs, while classification performance was largely at chance level for switch trials (except for the left dlPFC in adults and the object-selective ROI in children). Contrary to our expectations, we found no evidence that decoding accuracy differed between children and adults. In addition, task-set representations of the previously relevant task were not more likely to be (erroneously) decoded than the representations of the third task that was irrelevant on the previous and current trials. Thus, our analyses do not provide any evidence that task-set inertia contributed to lower distinctiveness for switch trials. The distinctiveness of the representation of the currently relevant task set decreased during switching in regions beyond the FP network, including CO regions associated with task control (Braver et al., 2003; Sestieri et al., 2014; Han et al., 2019; Palenciano et al., 2019; Cocuzza et al., 2020; Wood and Nee, 2023) as well as TO regions associated with the task-relevant stimuli (cf. Tsumura et al., 2021). Notably, children showed higher correlations in these decoding accuracy costs between FP and CO regions than adults, suggesting that switch demand affected task-set representations in FP and CO regions in a manner that was more similar among children than among adults.

### 4.1 More distinct representations on repeat than switch trials

The finding that the currently relevant task set could be decoded with an accuracy that was above chance for repeat but not for switch trials supports the notion that task-set representations in the present task were less stable when they had recently been updated (i.e., on switch trials; Meiran, 1996; Mayr and Kliegl, 2000). The lower distinctiveness on switch trials has partly been attributed to the lingering representation of the (no-longer relevant) task set from the previous trial (Rogers and Monsell, 1995; Wylie and Allport, 2000; Qiao et al., 2017; Rangel et al., 2023).

Directly testing this task-set inertia hypothesis in the present study, we did not find any evidence for this pattern, contrary to the previous findings reported by Qiao et al. (2017) based on representational similarity between consecutive trials. Note that compared to Qiao et al. (2017), our participants had to switch among three different tasks. This allowed us compare whether a multivariate pattern of brain activation contained more information of the previously relevant task than a third task (relevant on neither the previous nor the current trial), and thus directly test predictions made by the task-set inertia hypothesis.

Taken together with previous studies investigating neural representations during task switching (Loose et al., 2017; Qiao et al., 2017), the present results indicate that differences in neural task-set representations may depend on the specific kind of switching demand. Specifically, our paradigm and the one used by Qiao et al. (2017) required switches between different arbitrary rules and their corresponding response mappings that may be more likely to modulate the distinctiveness of task-set representations (cf. Woolgar et al., 2011), as indicated by a condition difference in decoding accuracy. In contrast, the paradigm by Loose et al. (2017) required switches between responses while the task remained the same conceptually, which resulted in comparable (above-chance) decoding for both switch and repeat trials (cf. Brass and De Baene, 2022).

### 4.2 Similar distinctiveness of task-set representations in children and adults

By demonstrating that the currently relevant task can be reliably predicted from neural activation patterns during task switching not only in adults but also in children, our results provide novel insights into children’s ability to flexibly switch between rules. These findings add to an emerging research direction investigating the role of neural representations for cognitive development across childhood and adolescence (e.g., Fandakova et al., 2019; Jung et al., 2023).

Contrary to our expectations based on comparisons of univariate task-based activation during task switching (e.g., Crone et al., 2006b; Wendelken et al., 2012), we did not find evidence for less distinct neural task-set representations in children compared to adults. However, a similar level of decoding accuracy between children and adults should not be taken as evidence that representations were identical or used in the same way in both age groups. Even with similar distinctiveness of task-set representations, there may still be differences in the application of the task set. Specifically, decoding techniques only indicate differences between task-set representations but do not reveal the processes through which these representations come to be or how they influence behavior (cf. Kriegeskorte and Douglas, 2018).

For example, tasks eliciting different levels of task performance in adults were shown to have comparable levels of decoding accuracy, suggesting that differences in complexity were not captured by the decoder (Ruge et al., 2019). This may be especially relevant when comparing groups of individuals of different ages, given that the neural systems that are being decoded differ in organization due to maturation and experience. In line with this consideration, Crone and colleagues (2006c) found that age differences in univariate activation were more pronounced when representations in working memory needed to be manipulated not just maintained. Finally, univariate analyses of the present sample revealed that children upregulated frontoparietal activation on switch compared to repeat trials to a smaller extent than adults did (Schwarze et al., 2023), further supporting the idea that children and adults may differ with respect to implementing the newly relevant task-set representation on switch trials.

### 4.3 Differences between networks

Corroborating and extending previous studies of neural task-set representations during rule-based tasks (Woolgar et al., 2011; Zhang et al., 2013; Loose et al., 2017; Qiao et al., 2017) with respect to regional heterogeneity, we showed greater decoding accuracy in TO regions compared to FP and CO regions, not only in children but also in adults. TO regions may be more strongly driven by the visual input of the task, support sensory representations within working memory (cf. Olivers and Roelfsema, 2020), or carry representations of lower dimensionality rendering them more distinct in their neural pattern (cf. Buschman, 2021). Note that the present task did not include multiple cues (i.e., a single cue was used for each task) and the cue was presented simultaneuously with the stimuli. We can thus not rule out that the cue contributed to the distinctiveness of task-set representations (cf. Loose et al., 2017). Given the relatively earlier position of the TO regions in the ventral visual processing stream (e.g., Kravitz et al., 2013), the extent to which the cue might have impacted task-set representations may differ between ROIs and thus contribute to differences in decoding accuracy between ROIs.

Our exploration of regional heterogeneity led to the finding that the difference between switch versus repeat decoding accuracy was correlated more strongly between the CO and FP ROIs (but not the TO ROIs) in children than in adults. It has been noted before that differences in the functional roles of CO and FP networks increase in the course of child and adolescent development (e.g., Fair et al., 2009; Keller et al., 2022; Tooley et al., 2022). In light of the individual differences observed here, we could speculate that the closer functional association between FP and CO regions in children is associated with the representation of more similar features in these regions in children as compared to adults. This idea is consistent with suggestions that representational structure is crucial for efficient cognitive control (Badre et al., 2021; Garner and Dux, 2023) and differs between regions in adults (Vaidya and Badre, 2022). A recent study examined how representational structures change during the acquisition of new tasks in adults (Mill and Cole, 2023) and demonstrated regional differences in compositional representations, that is, task-general activation patterns, and conjunct representations, that is task-specific activation patterns. Specifically, with learning, compositional representations in cortical regions were replaced by conjunct representations previously only found in subcortical regions.

### 4.3 Conclusion

Taken together, our results demonstrate that task-set representations were affected by switch demands – not only in adults but also in children. Individual differences in the degree to which effects of switch demands were correlated between sets of brain regions raise the possibility that the closer functional association between frontoparietal and cinguloopercular regions in children is related to the representation of more similar task features across regions in childhood. These findings raise further questions about the role of representations in the development of cognitive control during childhood that merit further study: What is the role of regional heterogeneity and overlapping feature representation for the learning and generalization of task rules in childhood? Future work focusing on developmental changes in neural representations could provide fruitful in elucidating the mechanisms underlying cognitive control development.

## Acknowledgments

We acknowledge financial support by the Max Planck Institute for Human Development and by the DFG Priority Program SPP 1772 “Human performance under multiple cognitive task requirements: From basic mechanisms to optimized task scheduling” (Grant No. FA 1196/2-1 to Y.F.). During the work on her dissertation S.A.S. was a predoctoral fellow of the International Max Planck Research School on the Life Course (LIFE; http://www.imprs-life.mpg.de) at the Max Planck Institute for Human Development, Berlin, Germany. The authors thank Theodoros Koustakas for support with the analyses. Communication regarding this article should be directed to S.A.S, or Y.F..

